# Application of the stochastic labeling methods with random-sequence barcodes for simultaneous quantification and sequencing of environmental 16S rRNA genes

**DOI:** 10.1101/072298

**Authors:** Tatsuhiko Hoshino, Fumio Inagaki

## Abstract

Next-generation sequencing (NGS) is a powerful tool for analyzing environmental DNA and provides the comprehensive molecular view of microbial communities. For obtaining the copy number of particular sequences in the NGS library, however, additional quantitative analysis as quantitative PCR (qPCR) or digital PCR (dPCR) is required. Furthermore, number of sequences in a sequence library does not always reflect the original copy number of a target gene because of biases caused by PCR amplification, making it difficult to convert the proportion of particular sequences in the NGS library to the copy number using the mass of input DNA. To address this issue, we applied stochastic labeling approach with random-tag sequences and developed a NGS-based quantification protocol, which enables simultaneous sequencing and quantification of the targeted DNA. This quantitative sequencing (qSeq) is initiated from single-primer extension (SPE) using a primer with random tag adjacent to the 5’ end of target-specific sequence. During SPE, each DNA molecule is stochastically labeled with the random tag. Subsequently, first-round PCR is conducted, specifically targeting the SPE product, followed by second-round PCR to index for NGS. The number of random tags is only determined during the SPE step and is therefore not affected by the two rounds of PCR that may introduce amplification biases. In the case of 16S rRNA genes, after NGS sequencing and taxonomic classification, the absolute number of target phylotypes 16S rRNA gene can be estimated by Poisson statistics by counting random tags incorporated at the end of sequence. To test the feasibility of this approach, the 16S rRNA gene of Sulfolobus tokodaii was subjected to qSeq, which resulted in accurate quantification of 5.0 × 10^3^ to 5.0 × 10^4^ copies of the 16S rRNA gene. Furthermore, qSeq was applied to mock microbial communities and environmental samples, and the results were comparable to those obtained using digital PCR and relative abundance based on a standard sequence library. We demonstrated that the qSeq protocol proposed here is advantageous for providing less-biased absolute copy numbers of each target DNA with NGS sequencing at one time. By this new experiment scheme in microbial ecology, microbial community compositions can be explored in more quantitative manner, thus expanding our knowledge of microbial ecosystems in natural environments.

## Introduction

Quantifying and characterizing the taxonomic composition and diversity of microbial communities in natural environments are primary foundations in microbial ecology. Quantitative PCR (qPCR) using DNA-binding fluorescent dyes [1] or sequence-specific probes (e.g., Taqman [2]) is a powerful and sensitive tool [3] for the quantification of a target gene, which has been widely used in environmental microbiology (e.g., 16S rRNA genes) and other biological research fields. However, these quantification methods use external standards and sometimes result in inaccurate values due to differences in the efficiency of PCR with “clean” standard DNA and “dirty” environmental DNA, which may also contain PCR-inhibiting substances [4–6]. The efficiency of PCR can also be affected by the GC content, secondary structure of the targeted sequence, bases adjacent to the 3’ end of the primers, and other factors [3, 7-15]. Those potential factors introducing biases always have some risks to produce accurate and hence reliable quantification results for the study of environmental microbial communities. Digital PCR (dPCR) is an approach that would circumvent the above-mentioned issues, because it is less affected by the PCR efficiency and provides the absolute copy number of DNAs without external standards [16, 17]. However, the both qPCR and dPCR quantification assay must be optimized for each target gene (or taxa), necessitating the design of specific primers and standardized PCR conditions on a taxon-by-taxon basis. Because the optimal condition (i.e. concentration of template DNA and annealing temperature) is different among different primers specific for a taxa. In general, such experimental processes are cumbersome and not likely amenable to high-throughput analyses.

NGS of PCR-amplified 16S rRNA genes has been used to study microbial community structures in a variety of environments, including the ocean [18, 19], soils [20, 21], and the human body [22, 23]. NGS enables the reading of tens of millions of sequences per run, permitting the analysis of even "rare biosphere" members of a microbial community that cannot be detected by conventional sequencing methods (e.g., Sanger method) [24, 25]. This advantage enables researchers to capture more comprehensive pictures of the naturally occurring microbial communities. For quantification of particular sequences in the NGS library, it is problematic that the proportion of sequence reads for each genetic component (e.g., phylotype in the case of 16S rRNA genes) in the sequence library is not directly linked to the number of target sequences in the template DNA due to differences in PCR efficiency for different target sequences [15, 26]. It has also been reported that different DNA polymerases and PCR conditions often resulted in different community structure determinations, indicating that comparative analysis between different laboratories based on NGS sequence read numbers is not straightforward and should be carefully considered [27, 28].

To address the issues described above, we employed the stochastic labeling approach [25] and optimized a protocol that enables the simultaneous determination of both 16S rRNA gene sequences and absolute copy number of the rRNA gene of particular phylotypes while avoiding the biases associated with PCR efficiency by introducing random-sequence barcodes before sequencing. The utilization of random-sequence barcodes has been reported for counting individual DNA molecule using microarray [29], and is has also been applied to sequencing for improving allele or mutant detection accuracy, correcting PCR errors causing artificial diversity [30–33]. Kivioja *et al.* used random-sequence barcodes during ligation before PCR amplification or during cDNA synthesis for genome scale human karyotyping and mRNA sequencing and quantification [34]. However, to the best of our knowledge, it has not been applied to absolute quantification with classification of environmental DNA in natural ecosystems. For absolute quantification of 16S rRNA gene, we introduced 65,536 random-sequence barcodes on the 5’ ends of primers used for single-primer extension (SPE), such that every target 16S rRNA gene would be stochastically labeled with a random tag. After the random barcode-labeled target DNAs were separated and sequenced using standard tag-sequencing methods, the absolute copy number of each target DNA sequence can be counted based on the incorporated random tags for each phylotype. Using this protocol, we could accurately quantify the 16S rRNA gene copy number of different phylotypes in environmental samples with sequence informations.

## Results and Discussions

### Method overview

Stochastic labeling of DNA molecules with a diverse array of random-sequence tags during the procedure of tag-sequencing was employed to quantify 16S rRNA genes in environmental DNA samples, allowing simultaneous sequencing and quantification. The quantitave sequencing (qSeq) procedure is summarized in Fig 1. The first step involves preparation of a primer for SPE that is composed of a sequence for indexing and sequencing, a random barcode tag (RBT: random octamer [N8] in this study), and a sequence complementary to that of the target genomic DNA in the 5’ to 3’ direction. This is followed by enzymatic digestion of excess SPE primers to avoid incorporation of RBT during subsequent PCR cycles. The SPE product is specifically amplified by PCR using primers targeting the SPE primer sequence and genomic DNA. Subsequently, indexing and adapter sequences are added to the PCR amplicon for Illumina sequencing. The resulting products contain the RBT sequences at the beginning of reads that are incorporated during the SPE step. Finally, the absolute number of target DNAs in the template environmental DNA sample is estimated from the number of RBT obtained.

**Fig 1.**
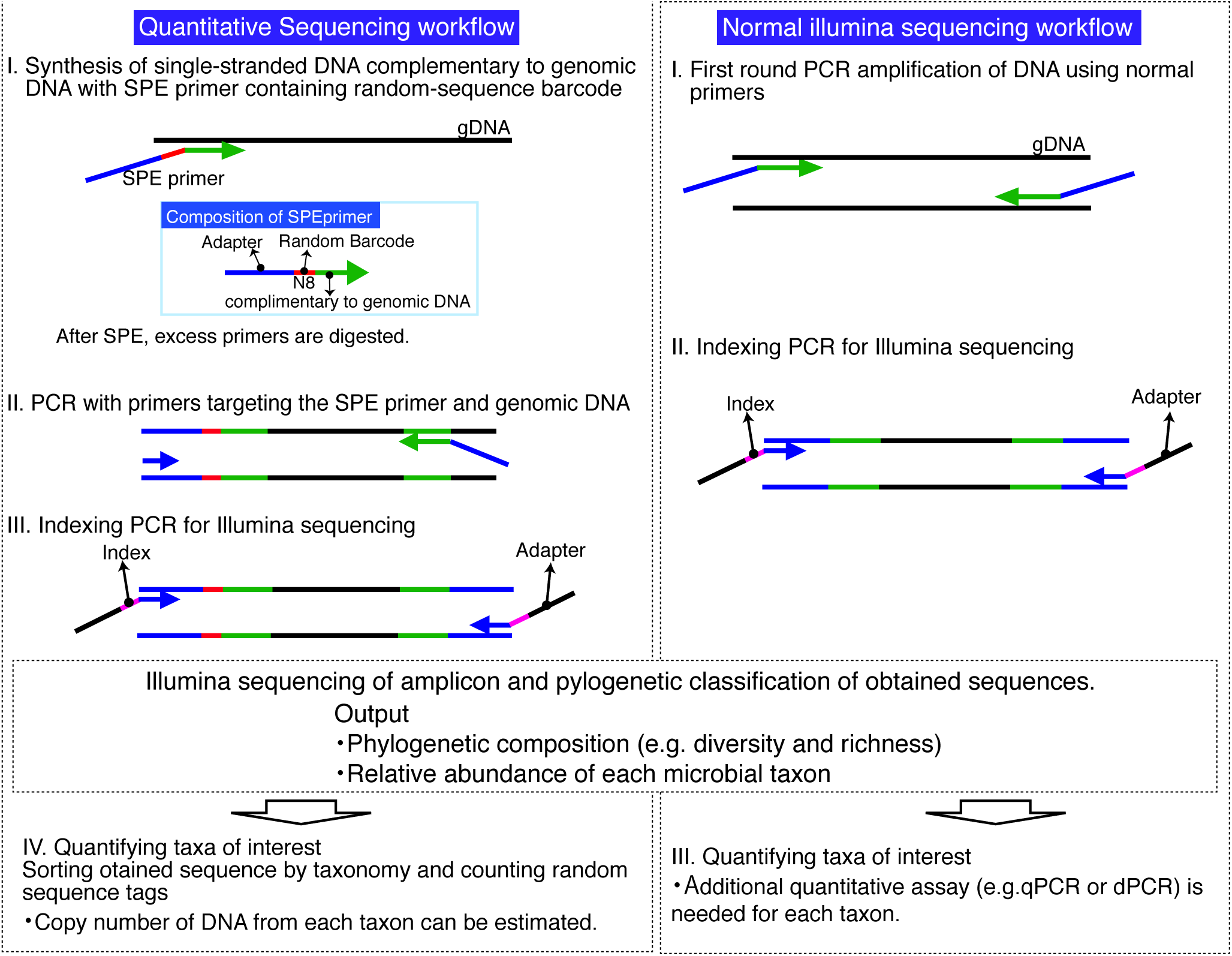
Schematic overview of the quantitative sequencing method and normal illumina sequencing workflows

This method has several advantages. First, counting the number of labels after sequence classification enables simultaneous quantification of many different sequences; e.g., at the level of bacterial or archaeal 16S rRNA genes or even finer levels of classification. Second, the quantification is independent of external DNA standards, as are used in qPCR, and because the number of random-barcode sequences is controlled by incorporation during the SPE reaction, it is not affected by the efficiency of subsequent PCR steps. Hence, quantification results are subject to less bias resulting from differences in PCR efficiency for different target DNAs and/or sampling sites.

The incorporation of the random-tag sequence into the SPE product is a stochastic process similar to dPCR [35, 36]; i.e., the number of containers in dPCR is the same as the number of random labels in the present method. The number of DNA molecules used as templates for SPE and subsequent PCR can be estimated based on a Poisson distribution. Thus, the number of target DNA molecules (N) is approximately given by equation 1, where C represents the the number of unique RBT (e.g., in this study, C is 4^8^ = 65,536, because random octamers [N8] is used for the RBT) and H represents the unique random tags incorporated into the SPE product, respectively:

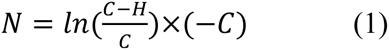

The expected increment (P) in number of observed unique RBT per every increment of a sequence at a given number (n) is determined using equation 2:

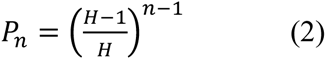

The observed number of unique RBT (S) is given by equation 3, because P_n_ is an arithmetic sequence where r = (H−1)/H:

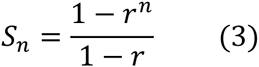

With in crease in n, S_n_ converges with H, in other word, by increasing the number of sequence reads, the number of unique RBT observed increases and gets close to the the number of incorporated unique random tags. Therefore, equation 4 is used to estimate the number of target DNA molecules when n is large enough relative to H:

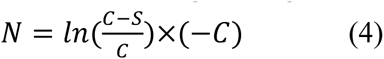

### Dynamic range test

Genomic DNA of *Sulfolobus tokodaii* with different concentrations from 5.0 × 10^3^ to 5.0 × 10^5^ copies were used to determine the dynamic range of the method. Precise quantification was achieved across the entire range of DNA copy numbers examined. Please note, however, that the accuracy at copy numbers over 1.0 × 10^5^ was relatively low (Fig 2), which may result in significant underestimation. This is most likely due to insufficient labeling because the number of labels is lower than the number of template DNA molecules. This relatively narrow dynamic range comparing to qPCR is a drawback of the current protocol, however the upper quantification limit could be increased simply by increasing the length of the random barcode; e.g., by adding 1 base to N8 would provide 4^9^ = 2.62 × 10^5^ barcode sequences. However, more than 10^6^ reads per sample are required to recover 10^5^ barcode sequences incorporated into the DNA. We anticipate that further developments in sequencing technologies will enable much higher throughput and resolve the trade-off issue, although applying right amount of template DNA below the upper limit would be straightforward. More accurate quantification (78-109% of expected) was observed in the DNA copy number ranging from 5.0 × 10^3^ to 5.0 × 10^4^. In this study, therefore, we utilized 5 × 10^3^ to 5 × 10^4^ copies of the target DNA for each qSeq analysis to maximize the quantification accuracy. We did not test fewer copy number than 5.0 × 10^3^ in this study because 5.0 × 10^3^ is roughly equal to ~5 pg of DNA and using that few DNA in NGS might not provide meaningful data upon 16S rRNA gene survey for natural microbial habitats. Slight underestimations observed with lower copy numbers could be caused, in part, by reduced reaction efficiency, especially the efficiency of SPE priming. To remedy this, we tested longer reaction times, higher primer concentrations, and slower ramp rates. Changing these parameters did not significantly improve the results, however. In future studies, improvement might be achieved using DNA analogues as locked nucleic acids, which have higher affinity for DNA in the SPE primer [37, 38].

**Fig 2.**
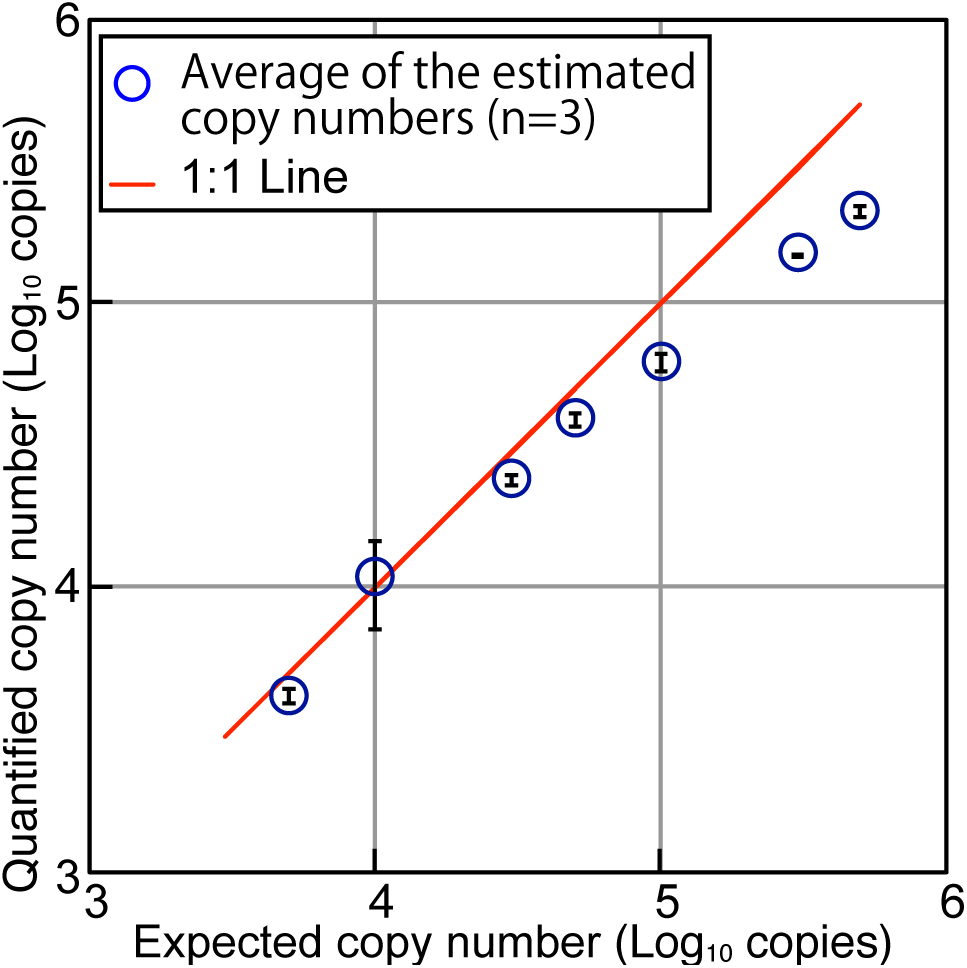
Plot of expected copy number versus copy number determined experimentally by qSeq. Genomic DNA of *S. tokodaii* was used for the test. Error bars indicate standard deviation from triplicate measurements.

Errors in PCR and/or sequencing represent another issue of concern, because the qSeq method counts any such error as a barcode sequence, potentially resulting in overestimation of target DNA molecules. To reduce the effect of these errors, the number of sequence reads used for the analysis should be kept to a minimum. In this study, we used 10-fold number of sequences for quantification compared with the expected copy number and did not observed such an overestimation in the dynamic range test.

### Analysis of mock microbial communities

A DNA solution containing identical copy numbers of the 16S rRNA genes of *Methanocaldococcus jannaschii*, *Halomonas elongata*, *S. tokodaii*, *Bacillus subtilis*, *Streptomyces avermitiis*, and *Paracoccus denitrificans* was sequenced by the standard Illumina sequencing and qSeq. The total copy number was 2.0 × 10^4^, which was within the dynamic range as described above. The number of RBT was counted after sorting of the qSeq results by phylogeny, and the 16S rRNA gene copy number was estimated using the equation as described above. As a result, qSeq could provide copy number of each microbial species and the quantified copy numbers were consistent with the expected value (Table 1). Based on the estimated copy number, relative abundances of each species were calculated (Fig 3A). In the standard sequencing analysis, the proportion of each species was calculated based on the relative ratio in the resulting sequence library. Standard sequencing library analysis underestimated the relative abundance of the 16S rRNA gene of *M. jannaschii*, whereas the *H. elongata* and *S. tokodaii* 16S rRNA were overestimated (Fig 3A). The GC content of these 16S rRNA genes, which is well known to affect PCR efficiency, is 31, 61, and 33% for *M. jannaschii, H. elongata* and *S. tokodaii*, respectively. Quantification bias cannot be predicted or explained based on GC content alone, and it is likely that other factors may be involved, such as priming efficiency, secondary structure, and/or bases adjacent to primers [7-10]. In contrast, the qSeq returned more accurate and reproducible data in good agreement with the expected ratios of the 16S rRNA gene copy numbers of each species examined.

**Table 1.** Quantification of 16S rRNA gene in the mock microbial community.

|  | Library | qSeq | Expected |
| --- | --- | --- | --- |
| Copy number (copies) |  |  |  |
| *S. tokodaii* | NA<sup>a</sup> | $(3.01 \pm 0.08) \times 10^3$ | $3.33 \times 10^3$ |
| *M. jannaschii* | NA | $(2.59 \pm 0.28) \times 10^3$ | $3.33 \times 10^3$ |
| *B. subtilis* | NA | $(2.89 \pm 0.16) \times 10^3$ | $3.33 \times 10^3$ |
| *S. avermitis* | NA | $(2.86 \pm 0.14) \times 10^3$ | $3.33 \times 10^3$ |
| *P. denitrificans* | NA | $(2.45 \pm 0.13) \times 10^3$ | $3.33 \times 10^3$ |
| *H. elongata* | NA | $(2.63 \pm 0.03) \times 10^3$ | $3.33 \times 10^3$ |

<sup>a</sup>NA indicates not applicable
Standard deviations are from triplicate measurements

In addition, two mock communities consisting of genomic DNA of two microbial species were tested. One was the mixture of the genomic DNA of *M. jannaschii* and *H. elongata* with 16S rRNA gene ratios ranging from 1:9 to 9:1 and the other was the mixture of *S. avermitilis* and *B. subtilis*. Both mock communities were subjected to the standard Illumina sequencing and the qSeq. The ratio of sequence reads in the sequence library was calculated and compared to the results obtained by the qSeq as described above. We found that 16S rRNA gene of *M. jannaschii* was always underestimated in the sequence library (t-test, p < 0.05 for all 3 plots), whereas the qSeq approach provided accurate ratios over the entire range tested (Fig 3B). This result was consistent with the results obtained from the 6-species mock community, indicating relative abundance in normal sequencing library can be biased as previously reported [10-13, 15]. We also tested the mock community of *S. avermitilis* and *B. subtilis* of which relative abundances in the 6-species mock community were similar (Fig 3A). As expected, both the ratios of *S. avermitilis* to *B. subtilis* obtained by sequence library and qSeq are consistent with expected ratio (Fig 3C).

**Fig 3.**
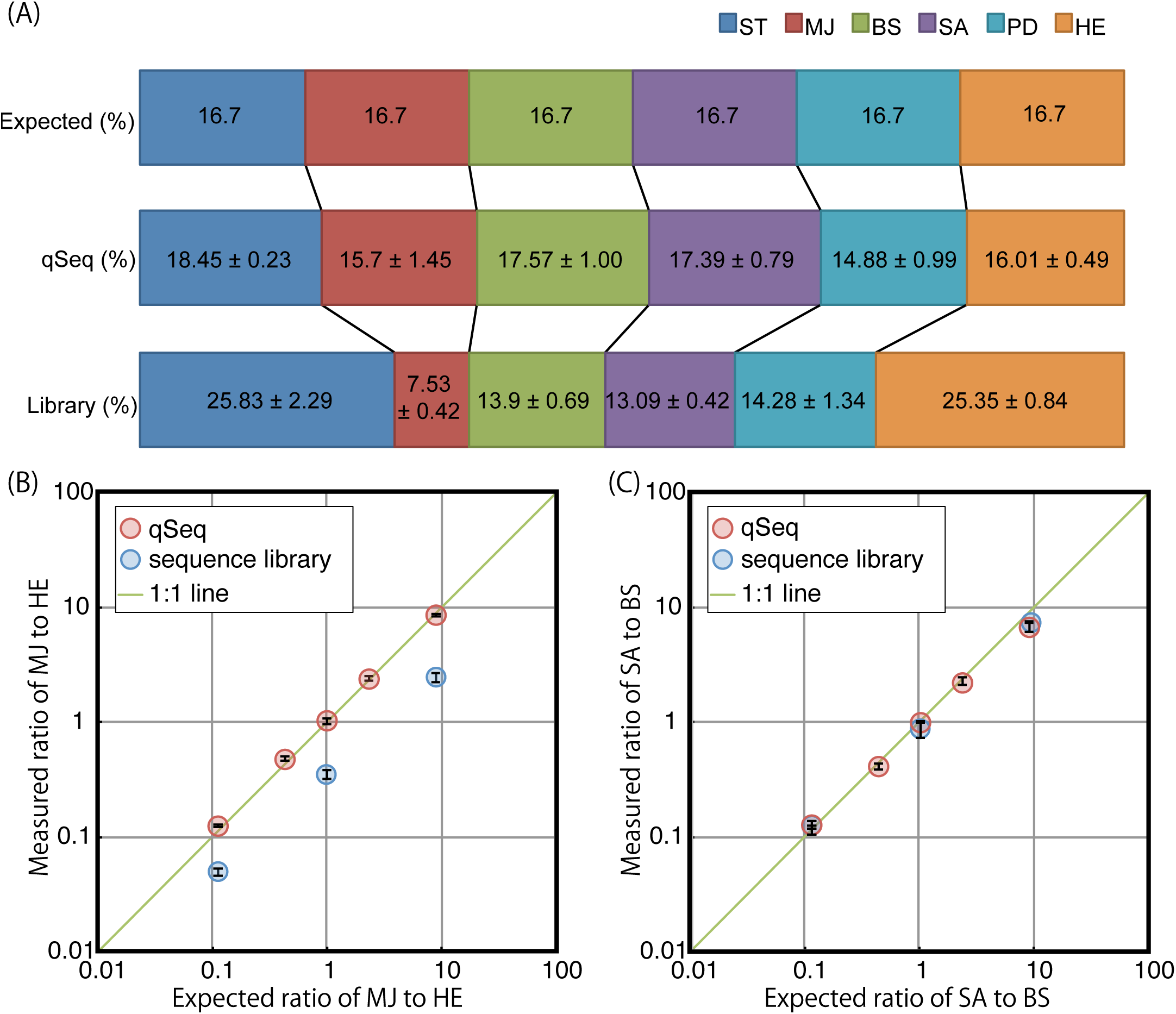
Analysis of mock microbial communities. (A) Mock microbial community of 6 micriboal species. Relative abundances were detemined by qSeq and standard illumina sequencing library. Standard deviation was calcurated from triplicate measurements. (B) Mock microbial community of 16S rRNA genes of HE and MJ at different ratios. Plot shows MJ to ME measured ratios versus expected ratio. (C) 16S rRNA genes of SA and BS at different ratios. Plot shows SA to BS measured ratios versus expected ratio. Error bars indicate standard deviation from triplicate measurements. ST, *S. tokodaii*; MJ, *M. jannaschii*; BS, *B. subtilis*; SA, *S. avermitiis*; PD, *P. denitrificans*; HE, *H. elongata*.

### Analysis of environmental samples

The qSeq approach was tested to some environmental samples, such as beach sand, paddy field soil, biofilm, and hot spring water sample. Bacterial and archaeal 16S rRNA genes were quantified separately for the reference using a microfluidic dPCR technique with domain-specific primers. The microfluidic dPCR enables determination of the absolute target DNA sequence copy number unaffected by biases introduced by differences in PCR efficiency [16]. Compared with the dPCR data, qSeq provided slightly lower copy numbers for bacterial and archaeal 16S rRNA genes in the paddy filed soil and biofilm sample (Fig 4A). In contrast, for the beach sand and hot spring water sample, qSeq provided higher copy numbers for bacterial and archaeal 16S rRNA genes. Although the results were not directly comparable due to differences in primer coverage, all the results for both techniques were within the same order of magnitude, indicating that qSeq enables accurate and precise quantification of those environmental DNA examined, at least as comparable to microfluidic dPCR. Moreover, the bacteria to archaea copy number ratios, which were determined using dPCR and qSeq as well as from the standard Illumina sequence library, were generally consistent: 0.60, 0.60, and 0.49, respectively, for biofilm; 0.43, 0.25, and 0.18, respectively, for hot spring; 0.024, 0.033, and 0.030, respectively, for paddy field soil; and 0.039, 0.040, and 0.060, respectively, for beach sand (Fig 4B). The ratios obtained from dPCR and qSeq were consistent each other in all the samples except for hot spring water, indicating, qSeq could quantify 16S rRNA without bias regarding amplification efficiency. In the hot spring sample, dPCR resulted in significantly higher ratio than the other two methods probably due to difference in coverage of the primer sets used for amplification. Overall, the result obtained from qSeq was shown to be as accurate as currently used methods for the analysis of environmental samples.

The qSeq protocol allows us to compare quantity of 16S rRNA gene in different environments. Table 2 shows an example for quantification in which the copy numbers of 16S rRNA gene of selected 5 classes were estimated by counting random sequence tags after classification of the sequence reads. *Methanomicrobia*, one of the classes containing methanogenic Archaea mainly from paddy and lakes, is detected in the paddy files soil to be (1.43±0.14) × 10^7^ copies/g-sample, whereas it is absent in the beach sand. As for Proteobacteria, the paddy field contains 5 times more β-Proteobacteria than the beach sand while ɣ-Proteobacteria populations were in the same order of magnitude. These examples demonstrates that qSeq has a great advantage that copy numbers can be directly quantified after sequencing without additional quantitative approaches as qPCR or dPCR.

**Fig 4.**
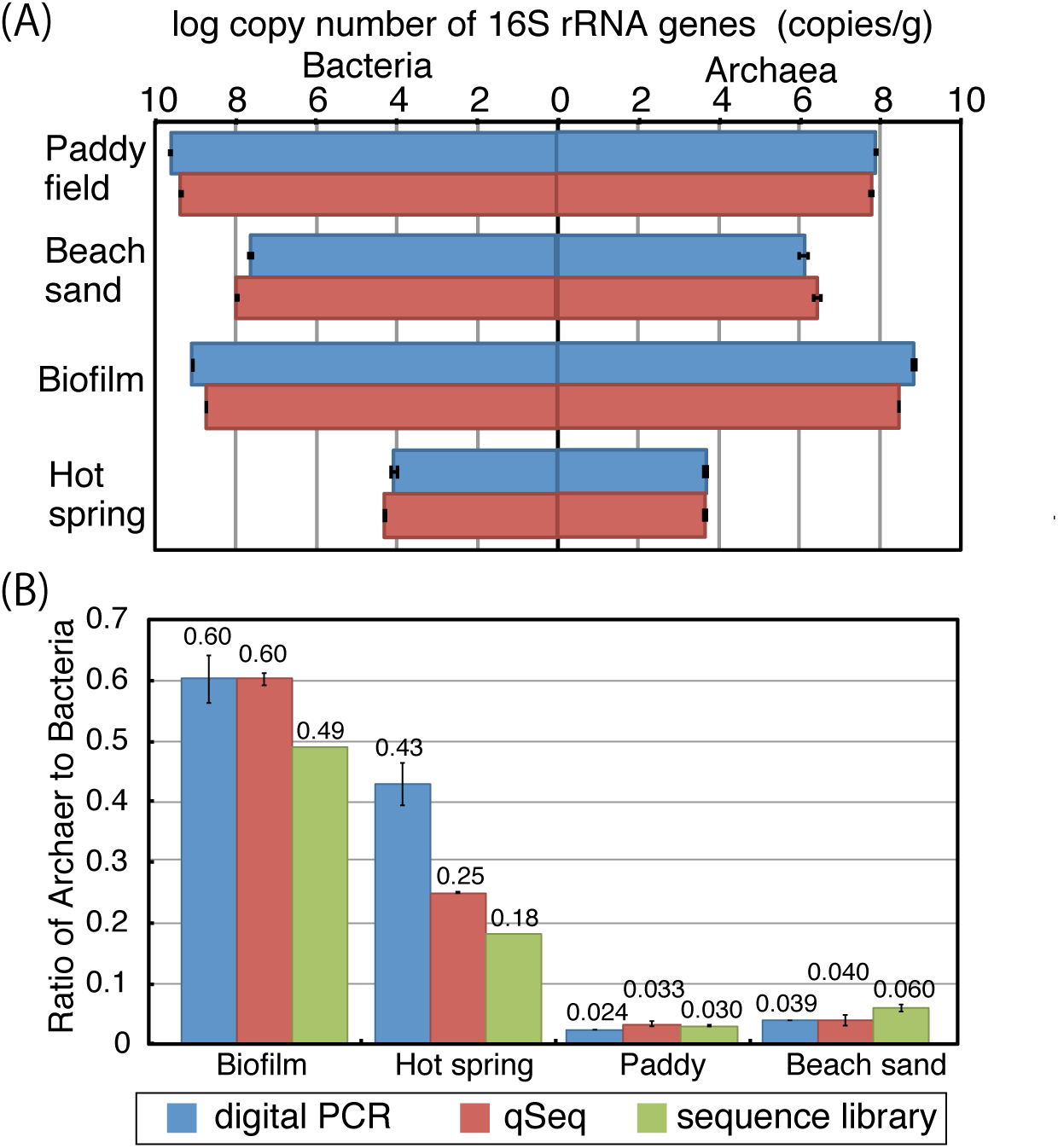
Analysis of environmental samples. (A) 16S rRNA genes were quantified by dPCR and qSeq using DNA extracted from the environmental samples. Error bars indicate standard deviation from triplicate measurements. (B) Ratio of archaea to bacteria determined by dPCR, qSeq and the standard Illumina sequencing library. Error bars indicate standard deviation from triplicate measurements. As for biofilm and hot spring sample, the ratio was calculated from single sample.

**Table 2.**
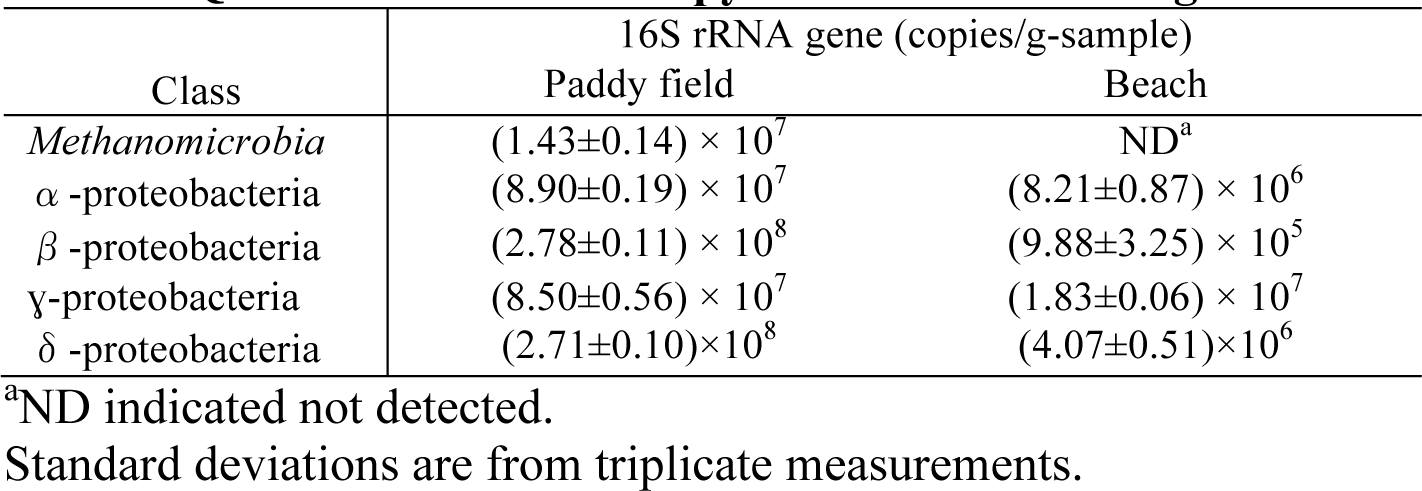
Quantification of the copy number 16S rRNA gene of selected classes.

| Class | 16S rRNA gene (copies/g-sample) |  |
| --- | --- | --- |
|  | Paddy field | Beach |
| *Methanomicrobia* | $(1.43 \pm 0.14) \times 10^7$ | ND<sup>a</sup> |
| α-proteobacteria | $(8.90 \pm 0.19) \times 10^7$ | $(8.21 \pm 0.87) \times 10^6$ |
| β-proteobacteria | $(2.78 \pm 0.11) \times 10^8$ | $(9.88 \pm 3.25) \times 10^5$ |
| γ-proteobacteria | $(8.50 \pm 0.56) \times 10^7$ | $(1.83 \pm 0.06) \times 10^7$ |
| δ-proteobacteria | $(2.71 \pm 0.10) \times 10^8$ | $(4.07 \pm 0.51) \times 10^6$ |

<sup>a</sup>ND indicated not detected.
Standard deviations are from triplicate measurements.

## Conclusion

In summary, in analyses of a mock microbial community and actual environmental samples, we demonstrate that the qSeq protocol optimized for environmental DNA enables simultaneous sequencing and gene quantification. The absolute copy number of *S. tokodaii* 16S rRNA gene was accurately quantified by qSeq in the template copy range of 10^3^ to 5 × 10^4^. The mock microbial community tests showed that qSeq enables accurate determinations of the ratios of microbial community members and that the technique would be less affected by possible biases due to differences in PCR efficiency. The quantification accuracy and precision of bacterial and archaeal 16S rRNA genes in environmental samples assessed by qSeq were comparable to microfluidic dPCR. The qSeq technique described here enables parallel sequencing and quantification of many different target DNAs at one time, and we anticipate that it will be of great value in future studies in environmental microbiology.

## Materials and Methods

### Sample collection and DNA extraction

Beach sand and soil samples were collected from the coast and from a paddy field near the Kochi Institute for Core Sample Research, JAMSTEC, in Nankoku, Kochi, Japan, using a 50 ml tip-cut syringe for the each site. Biofilm sample and hot spring water sample were collected at a stream in Garandake which is an active volcano located in Oita, Japan. The collected hot spring water was filtrated by a polycarbonate filter at the sampling site.

All the sample including the filter were stored at −80°C prior to analysis. DNA was extracted using a PowerMax Soil DNA Isolation Kit (MOBIO). The extracted DNA was further concentrated by isopropanol precipitation and finally resuspended in TE buffer.

### Genomic DNA and mock community preparation

Genomic DNA (gDNA) of *Paracoccus denitrificans* (catalog no. JGD12662), *Sulfolobus tokodaii* (catalog no. JGD12662), *Halomonas elongata* (catalog no. JGD08103), *Methanocaldococcus jannaschii* (catalog no. JGD12154), *Bacillus subtilis* subsp. *subtilis* (catalog no. JGD08099), and *Streptomyces avermitilis* (catalog no. JGD12254) were provided by the RIKEN DNA Bank through the National Bio-Resource Project of the MEXT, Japan. DNA concentration was determined using a Nanodrop 3300 fluorospectrometer with picogreen staining (Invitrogen). The copy number of 16S rRNA gene on each gDNA were determined using whole genome sequences found in the GenBank database. Based on the measured DNA concentration, the copy number of 16S rRNA on gDNA, and molecular weight of each gDNA, copy number of 16S rRNA per pg of DNA was calculated. *S.tokodaii* gDNA was subsequently diluted with EASY Dilution buffer (TaKaRa Bio) to the desired concentration for the dinamic range test. For preparing the mock microbial communities, each gDNA from each species was at first diluted to 10^4^ copies of 16S rRNA gene/µl. Subsequently, the diluted gDNAs were mixed to obtain desired composition of mock microbial community.

### Standard Illumina DNA sequencing

To compare relative quantification results for library sequences using random-labeling quantification, standard Illumina sequencing was performed as described elsewhere [39, 40], using the universal primers N8-515F and N8-806R targeting the V4 region of 16S rRNA genes (Table 1). After purification of the PCR product of the expected size, indexing PCR was performed using a Nextera XT indexing kit (Illumina), followed by sequencing using a MiSeq platform with MiSeq Reagent Kit v2 for 500 cycles or v3 for 600 cycles (Illumina).

### Quantification by qSeq

For qSeq, SPE was performed using MightyAmp DNA polymerase ver. 2 (TaKaRa Bio) with 0.3 µM N8-515F primer in a total reaction volume of 20 µl. The amount of template DNA was 300 pg for environmental samples, and 2.0 × 10^4^ copies of 16S rRNA gene were used for the analysis of a mock microbial community sample. For dynamic range testing, 5.0 × 10^3^ to 5.0 × 10^5^ copies of 16S rRNA gene from the genomic DNA of *S. tokodaii* were used. The SPE conditions consisted of initial denaturation at 98°C for 2 min, cooling to 68°C at 0.3°C/s, and elongation at 68°C for 10 min. Subsequently, excess primers were digested by addition of 4 of Exonuclease I (5 U/µl, TaKaRa Bio) and incubation at 37°C for 120 min, followed by inactivation of the enzyme by incubation at 80°C for 30 min. A 5-µl volume of the SPE product was used for the first-round PCR; each 20-µl reaction contained 0.3 µM N8-U806R and F2-primers (Table 3), 1× MightyAmp buffer ver. 2, and 0.4 µl of MightyAmp DNA polymerase. Thermal cycling consisted of initial denaturation at 98°C for 2 min, followed by 40 cycles of 98°C for 10 s, 55°C for 15 s, and 68°C for 30 s. Finally, 15 µl of each PCR product was subjected to agarose gel electrophoresis, and the band of the expected size was excised and purified using a Nucleospin column (TaKaRa Bio). The purified PCR products were finally subjected to indexing PCR and sequencing by Miseq, as described above in the section of Standard Illumina DNA sequencing.

**Table 3.**
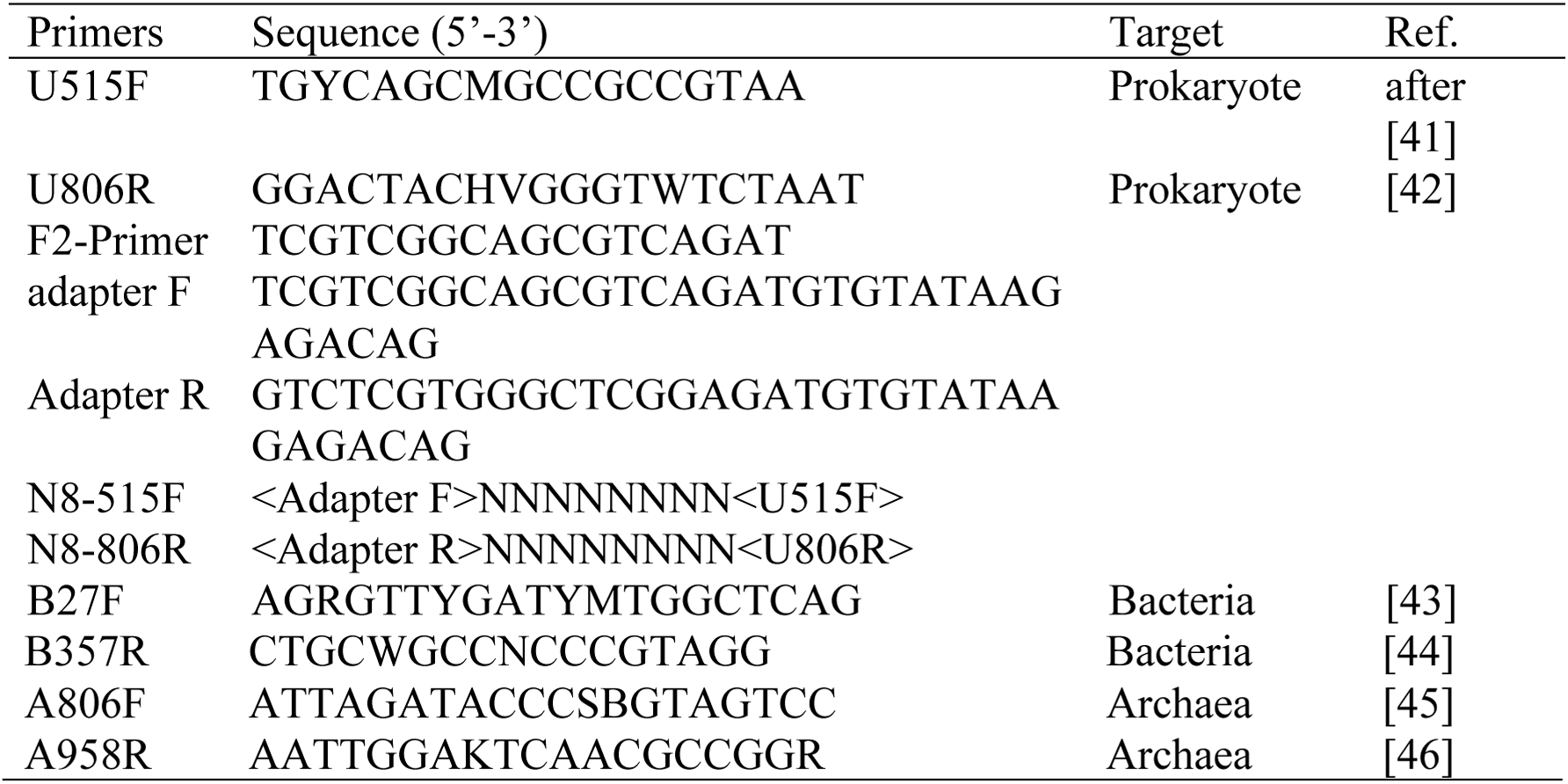
Sequences of oligonucleotide primers used in this study.

### Digital PCR

The absolute copy numbers of bacterial and archaeal 16S rRNA genes in environmental samples were determined by microfluidic dPCR using a BioMark Real-time system and 12.765 Digital Array (Fludigm) as described previously [16]. The domain-specific primer pairs B27F-B357R and A806F-A958R were used for bacterial and archaeal 16S rRNA genes, respectively (Table 3).

### Data analysis

All sequence data obtained in the study were processed using the Mothur software package (v.1.35.0) [47] for counting the unique random barcodes and for phylogenetic identification of the sequence reads. For qSeq analysis, 5.0 × 10^4^ and 2.0 × 10^5^ sequence reads were analyzed for quantification of the mock microbial communities and the environmental samples, respectively. At first, the sequence reads generated by illumina Miseq were assembled, trimmed and filtered using Mothur software package: screen.seqs and pcr.seqs with the sequences of U515F and U806R primers were used for screening sequences by length and primer match, respectively. Subsequently, classify.seqs with silva.nr_v123.align as a reference database was used for classification of rRNA gene. Based on the classification, get.lineage command sorted sequences of taxa of interest. RBT sequences were obtained using pcr.seqs with the reverse complementary sequence of U515F. After screening the obtained sequences by length, the number of RBT was determined using unique.seqs command.

### Deposit of DNA sequences in databases

The sequence data were deposited in the DDBJ/EMBL/GenBank databases under accession numbers DRA004531, DRA004792 and DRA004912.

### Ethics statement

The beach sample was collected from public beach area and the approved by Kochi prefectural office. Sampling from the paddy field was permitted by Kochi University. The hot spring water and biofilm samples were collected with the permission of Beppu-Hakudokogyo lnc.

## Acknowledgements

The authors are grateful for the technical assistance of S. Hashimoto, Y. Hamada, and T. Terada. The authors are also acknowledge M. Tanaka and M. Murayama for the assistance in sampling. This work is supported by the Japan Society for the Promotion of Science (JSPS) Grand-in-Aid for Science Research (no. 26251041 to FI, no. 15K14907 to TH).

